# Wnt7a Suppresses Adipogenesis of Skeletal Muscle Mesenchymal Stem Cells and Fatty Infiltration Through the Alternative Wnt-Rho-YAP/TAZ Signaling Axis

**DOI:** 10.1101/2022.06.30.498285

**Authors:** Chengcheng Fu, Mariana Guzmán-Seda, Damien Laudier, Woojin M. Han

## Abstract

Intramuscular fatty infiltration in muscle injuries and diseases, caused by aberrant adipogenesis of fibro-adipogenic progenitors, negatively impacts function. Intramuscular delivery of Wnt7a offers a promising strategy to stimulate muscle regeneration but its effects on adipogenic conversion of fibro-adipogenic progenitors remain unknown. Here we show that Wnt7a inhibits adipogenesis of FAPs through the alternative Wnt-Rho-YAP/TAZ signaling that subsequently upregulates the canonical Wnt pathway. Furthermore, intramuscular injection of Wnt7a in vivo effectively suppresses fatty infiltration in mice. Our results collectively suggest Wnt7a as a potential protein-based therapeutic for inhibiting adipogenesis of FAPs and intramuscular fatty infiltration in pathological muscle injuries or diseases.

## INTRODUCTION

Persistent fatty infiltration is a common hallmark of chronic skeletal muscle injuries and diseases that negatively impacts function and poses significant health and socioeconomic burden (Minagawa et al., 2013; Yamamoto et al., 2010). For example, in rotator cuff injuries, irreversible fatty infiltration is prevalent in the associated muscles, which directly increases muscle dysfunction and retear rates following surgical repair (Fu et al., 2021; Gladstone et al., 2007; Park et al., 2015b). In muscular dystrophies, pervasive intramuscular fatty infiltration positively correlates with the disease severity (Li et al., 2015). Fatty infiltration is also common in the paraspinal and neck muscles of astronauts following spaceflights (Burkhart et al., 2019; McNamara et al., 2019). Recent evidence suggests that fibro-adipogenic progenitors (FAPs), a population of muscle-resident mesenchymal stromal cells, are the primary cellular culprit that generates intramuscular fatty infiltration (Joe et al., 2010; Liu et al., 2016; Uezumi et al., 2010; Wosczyna et al., 2012). Although this link between FAPs and fatty infiltration is established, therapies that limit such pathologic fatty infiltration without compromising myogenesis currently do not exist.

FAPs play a critical role in muscle regeneration by secreting paracrine factors that promote activation and expansion of muscle stem/satellite cells (Joe et al., 2010; Uezumi et al., 2010; Wosczyna et al., 2012). Upon muscle injury, immune cells initially infiltrate the injured space to remove debris and activate both FAPs and satellite cells (Butterfield et al., 2006; Heredia et al., 2013; Tidball and Villalta, 2010). As the inflammation resolves, TNFα released by macrophages induces apoptotic clearances of FAPs while activated satellite cells continue to undergo myogenesis (Lemos et al., 2015). In chronic muscle pathology, however, FAPs undergo unchecked proliferation and give rise to adipocytes and myofibroblasts (Lemos et al., 2015). The resulting fatty infiltration and fibrosis perturb the highly aligned and organized muscle structure and consequently reduce the ability of muscles to contract and regenerate. Thus, identifying molecular mechanisms that regulate the adipogenic conversion of FAPs is critical for establishing strategies to combat pathologic fatty infiltration.

Intramuscular delivery of Wingless-type MMTV Integration Site Family 7a (Wnt7a) offers a promising strategy to both stimulate muscle regeneration and prevent muscle degeneration (Han et al., 2019; von Maltzahn et al., 2012, 2013; Schmidt et al., 2020). Wnt7a promotes myofiber hypertrophy through the non-canonical Akt/mTOR pathway and increases the symmetric expansion of muscle satellite cells through the non-canonical planar cell polarity pathway (Le Grand et al., 2009; von Maltzahn et al., 2011). In pre-clinical models of Duchenne muscular dystrophy, Wnt7a administration significantly increases satellite cell quantity, myofiber hypertrophy, and muscle strength (von Maltzahn et al., 2012). Controlled delivery of Wnt7a using a bioengineered hydrogel also increases satellite cell quantity and myofiber hypertrophy, presenting Wnt7a as an effective therapeutic candidate for treating various acute and degenerative muscle conditions (Han et al., 2019). While the past findings collectively corroborate that Wnt7a can be used as a potential pro-myogenic therapeutic, the effect of Wnt7a on FAPs, and specifically whether it limits adipogenic conversion of FAPs, remains unknown.

The objective of this study was to determine the mechanistic effect of Wnt7a on FAPs adipogenic conversion. By using freshly isolated primary murine FAPs, we demonstrate that Wnt7a effectively inhibits its adipogenesis through the alternative Rho-YAP/TAZ pathway. While Wnt7a does not directly increase nuclear localization of β-catenin, Wnt7a-induced nuclear localization of YAP/TAZ increases the production of canonical Wnt signaling modulators that further suppress adipogenesis. Importantly, Wnt7a treated FAPs adipogenic conditions do not adopt a fibroblastic phenotype. Finally, we also show that Wnt7a inhibits intramuscular fatty infiltration upon glycerol injury without negatively impacting myogenesis nor causing fibrosis in vivo. Collectively, we provide mechanistic evidence that Wnt7a effectively inhibits adipogenesis of FAPs and intramuscular fatty infiltration.

## RESULTS

### Wnt7a inhibits adipogenesis of FAPs

To evaluate the effect of Wnt7a on lineage specification of differentiating fibro-adipogenic progenitors (FAPs), we carried out a series of in vitro experiments using primary murine FAPs. Primary FAPs (CD31-, CD45-, α7 integrin-, Sca-1+) were isolated from the hindlimb muscles of C57Bl6/J mice using previously reported methods (**Fig. 1A**) (Marinkovic et al., 2019). Freshly isolated FAPs express PDGFRα (**Fig. 1B**), a defining marker of murine FAPs (Joe et al., 2010; Uezumi et al., 2010). Primary FAPs uniformly differentiated into myofibroblasts characterized by stress fibers expressing α-smooth muscle actin (αSMA) when maintained in fibrogenic media (**Fig. 1C, D**; p<0.001); but when maintained in adipogenic media, FAPs homogeneously differentiated into oil red O (ORO)+/perilipin-1+ adipocytes (**Fig. 1C-F**; p<0.001). Collectively, these results confirm the functional identity of the isolated primary FAPs to be used in the subsequent experiments.

**Figure 1.**
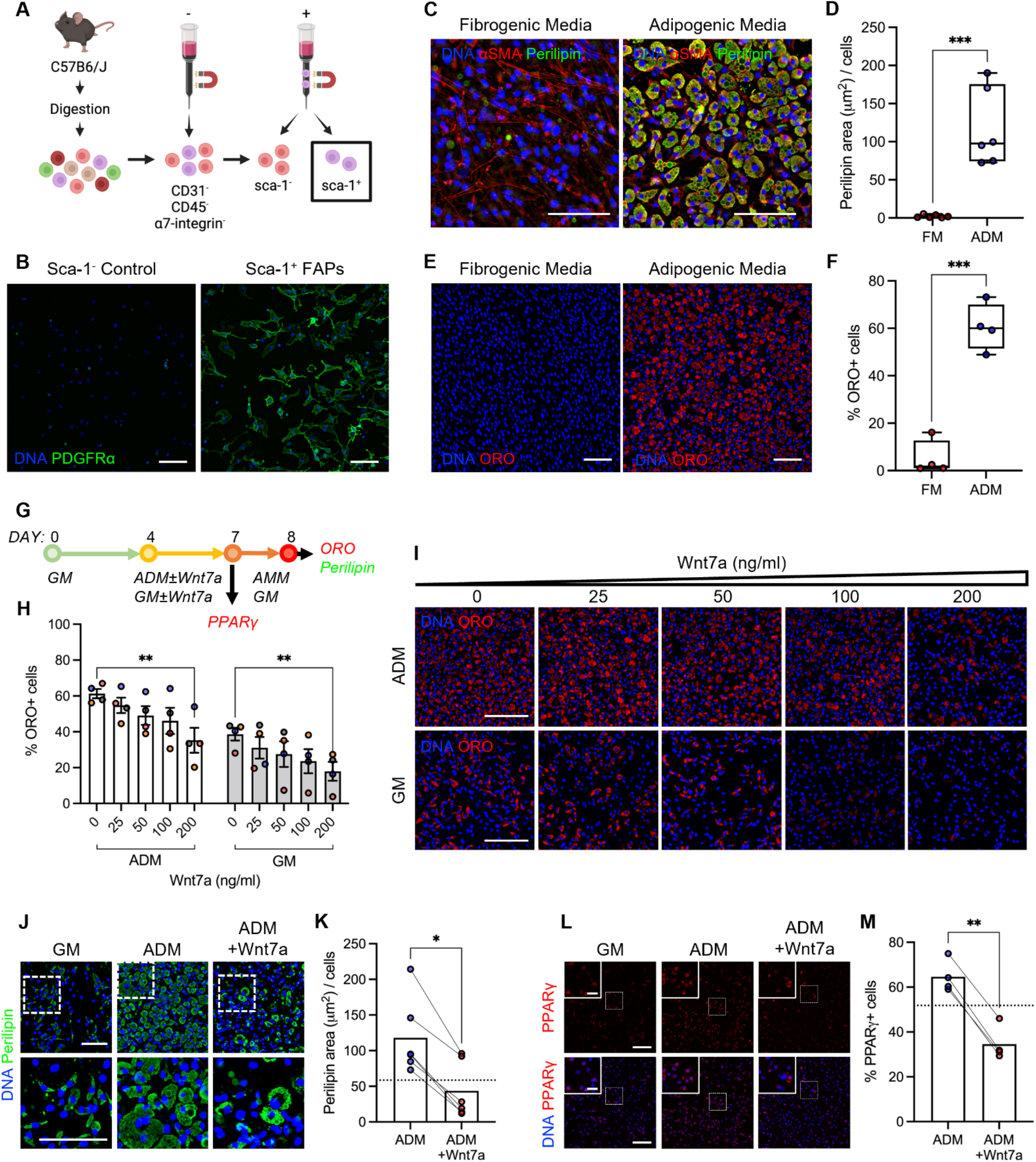
Wnt7a inhibits FAPs adipogenesis. ***(A)*** Schematic of FAPs isolation from C57BL6/J hindlimb muscles via magnetic activated cell sorting. Created with Biorender.com. ***(B)*** Freshly isolated FAPs express PDGFRα. 24 hours post-seeding. Scale bar: 100 µm. ***(C)*** Representative immunofluorescence images of α-smooth muscle actin (αSMA) and perilipin-labeled FAPs differentiated in fibrogenic and adipogenic differentiation media. Scale bar: 100 µm. ***(D)*** Perilipin area normalized by cell quantity in fibrogenic (FM) and adipogenic (ADM) media. Unpaired t-test. *** *p*<0.001. n=6. ***(E)*** Representative images of Oil Red O (ORO)-labeled FAPs differentiated in fibrogenic and adipogenic differentiation media. ***(F)*** Percent ORO+ cells in fibrogenic (FM) and adipogenic (ADM) media. Unpaired t-test. *** *p*<0.001. n=4. ***(G)*** Experimental timeline of Wnt7a study. GM: growth media; ADM: adipogenic differentiation media; AMM: adipogenic maintenance media. ***(H)*** Percent ORO+ cells treated with varying doses of Wnt7a. 2-way ANOVA with Bonferroni post-hoc analyses. Mean ± SEM. Dose effect p=0.0021. Media effect p<0.0001. ** *p*<0.01. n=4. Colors represent biological replicates. ***(I)*** Representative images of ORO-labeled cells treated with varying doses of Wnt7a. Scale bar: 100 µm. ***(J)*** Representative immunofluorescence images of perilipin-labeled FAPs cultured in GM and ADM ± Wnt7a (200 ng/ml). Scale bar: 100 µm. ***(K)*** Perilipin area normalized by cell quantity. Unpaired t-test. * *p*<0.05. n=6. Dotted line: Mean of GM. ***(L)*** Representative immunofluorescence images of PPARγ-labeled FAPs cultured in GM and ADM ± Wnt7a (200 ng/ml). Scale bar: 100 µm. Inset scale bar: 25 µm. ***(M)*** Percent PPARγ+ nuclei. Unpaired t-test. ** *p*<0.01. n=4. Dotted line: Mean of GM.

To determine the dose-dependent effect of Wnt7a on the adipogenic potential of FAPs, we seeded freshly isolated FAPs in a dish and let the cells proliferate to near confluency for 4 days in growth media. Proliferating FAPs were further maintained in growth media (±Wnt7a) or adipogenic differentiation media (±Wnt7a) for additional 3-4 days (**Fig. 1G**). Note, in growth media, the FAPs begin to spontaneously differentiate into both myofibroblasts and adipocytes (Joe et al., 2010). In both growth and adipogenic conditions, Wnt7a decreased adipogenesis in a dose-dependent manner (**Fig. 1H, I**). The dose of 200 ng/ml significantly reduced the formation of ORO+ adipocytes compared to the control (0 ng/ml; **Fig. 1H, I**; p<0.01). Based on this, we chose to use 200 ng/ml for the subsequent in vitro experiments.

To further validate the effect of Wnt7a on the suppression of FAPs adipogenesis, we cultured freshly isolated FAPs to near confluency and subsequently maintained the cells in either growth and adipogenic differentiation media, with or without 200 ng/ml Wnt7a (**Fig. 1G**). Note, the dH_2_O vehicle for Wnt7a (0.2% v/v in media) does not affect FAPs adipogenesis (**Supp. Fig. 1**). Wnt7a significantly reduced the formation of perilipin-1+ (lipid droplet-associated protein) adipocytes (**Fig. 1J,K**; p<0.0001) as well as nuclear activation of PPARγ, a master regulator of adipogenesis, compared to the control (**Fig. 1L,M**; p<0.01). In the adipogenic condition, Wnt7a also reduced perilipin-1 and PPARγ expressions to below the mean of spontaneously differentiating condition (**Fig. 1J-M**). Importantly, Wnt7a did not increase the fibrogenic potential of FAPs (**Supp. Fig. 2A**). In the spontaneously differentiating condition, the co-emergence of perilipin-1+ adipocytes and αSMA+ stress fiber-expressing myofibroblasts was apparent (**Supp. Fig. 2A**). However, treating FAPs with Wnt7a in the adipogenic condition did not promote stress fiber expressing myofibroblasts, indicating that Wnt7a does not promote fibrogenesis in vitro (**Supp. Fig. 2A**). Wnt7a treatment also did not affect cell viability (**Supp. Fig. 2B,C**). Altogether, these results demonstrate that Wnt7a inhibits adipogenesis of FAPs in a dose-dependent manner.

### Wnt7a inhibits adipogenesis of FAPs through non-canonical Wnt pathways

The canonical Wnt signaling inhibits adipogenesis through suppression of the adipogenic transcription factor PPARγ in preadipocytes, marrow-derived mesenchymal stromal cells, and muscle FAPs (Bennett et al., 2002; Kang et al., 2007; Longo et al., 2004; Moldes et al., 2003; Reggio et al., 2020; Ross et al., 2000). To determine if the Wnt7a isoform inhibits adipogenesis of FAPs through the canonical Wnt pathway, we cultured freshly isolated FAPs in the growth media for 4 days, then subsequently treated the cells in the adipogenic differentiation media containing Wnt7a (200 ng/ml) or the prototypically canonical activator Wnt3a (200 ng/ml) for 4 hours (**Fig. 2A**). FAPs treated in the adipogenic differentiation media containing dH_2_O vehicle and Wnt7a did not exhibit nuclear localization of β-catenin (**Fig. 2B,C**; >1000 cells assayed per condition from 3 biological replicates; **Supp. Fig. 3A**). However, brief 4-hour treatment with Wnt3a exhibited significantly increased nuclear localization of β-catenin compared to both control and Wnt7a conditions (**Fig. 2B,C**; p<0.05 vs Wnt7a, p<0.01 vs control). This suggests that Wnt7a activates β-catenin-independent and non-canonical Wnt pathways in primary murine FAPs.

**Figure 2.**
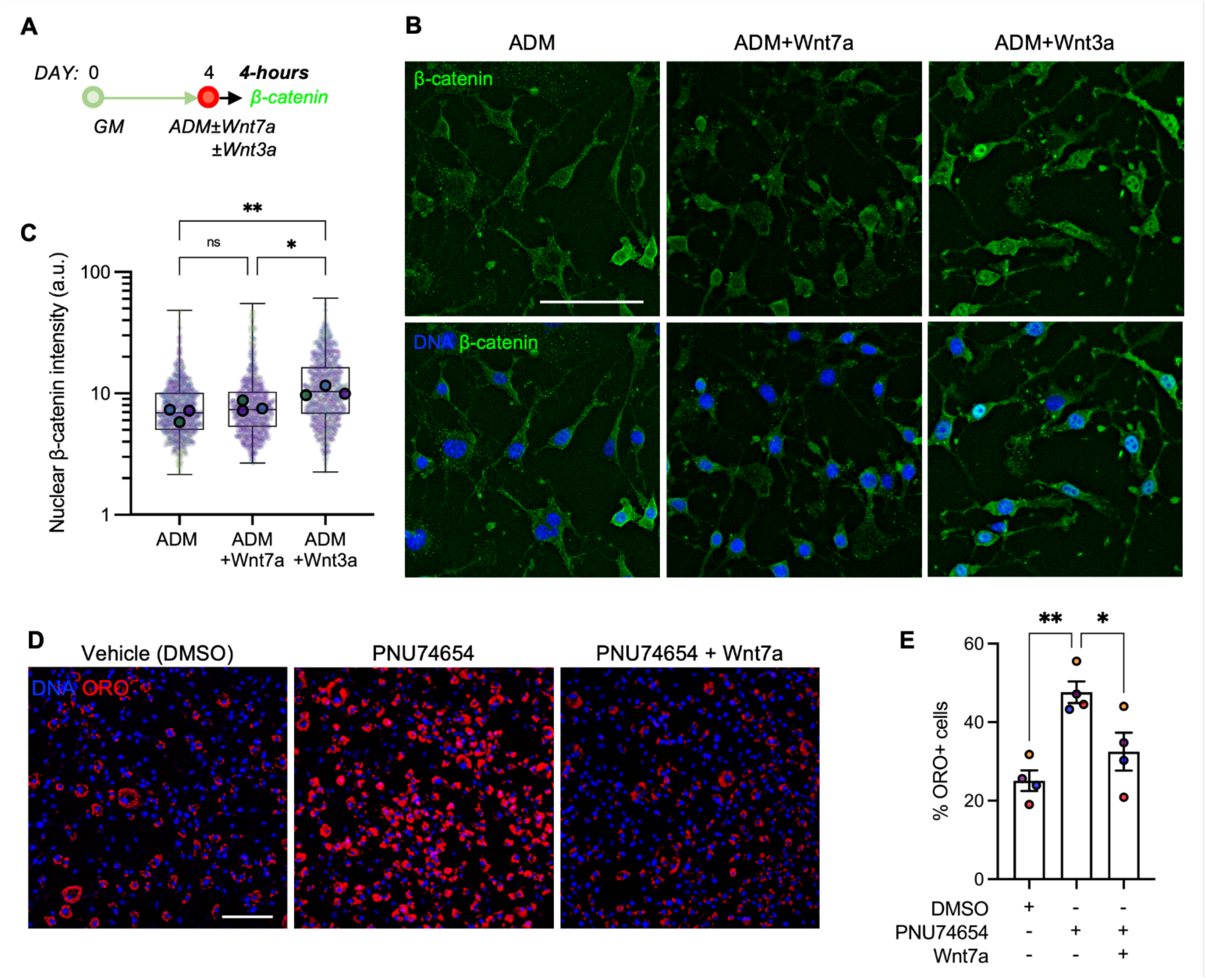
Wnt7a inhibits FAPs adipogenesis through non-canonical pathways. ***(A)*** Experimental timeline of β-catenin expression study. GM: growth media; ADM: adipogenic differentiation media. ***(B)*** Representative immunofluorescence images of β-catenin-labeled FAPs treated with vehicle, Wnt3a (200 ng/ml) and Wnt7a (200 ng/ml). Scale bar: 100 µm. ***(C)*** Nuclear β-catenin intensity. >1000 cells analyzed per condition in an automated manner from n=3 biological replicates. 1-way ANOVA with Tukey’s post-hoc analyses applied on the medians of biological donors. * *p*<0.05; ** *p*<0.01. n=3. Colors represent biological replicates. ***(D)*** Representative images of ORO-labeled cells treated with DMSO, PNU74654 (50 µM), PNU74654 (50 µM) + Wnt7a (200 ng/ml). Scale bar: 100 µm. ***(E)*** Percent ORO+ cells with DMSO, PNU74654, and PNU74654 + Wnt7a treated conditions. 1-way ANOVA with Tukey’s post-hoc analyses. Mean ± SEM. * *p*<0.05; ** *p*<0.01. n=4. Colors represent biological replicates.

To further corroborate this finding, we next sought to inhibit the activity of β-catenin using a small molecule inhibitor, PNU-74654. This inhibitor specifically prevents the binding of β-catenin and the T-cell factor, Tcf, in the nucleus. In this assay, we found that concentrations beyond 50 μM diminish FAPs proliferation in vitro, and thus we chose to use 50 μM for the inhibition study (**Supp. Fig. 3B**). As expected, inhibiting β-catenin/Tcf binding with PNU-74654 significantly increased FAPs adipogenesis compared to the DMSO vehicle control (**Fig. 2D,E**; p<0.01). However, co-treating FAPs with Wnt7a and PNU-74654 resulted in a marked reduction in adipogenesis compared to the PNU-74654 condition (**Fig. 2D,E**; p<0.05). Because Wnt7a does not increase nuclear expression of β-catenin (**Fig. 2B**) but reduces adipogenesis while β-catenin remains inhibited (**Fig. 2D**), Wnt7a may be acting through a non-canonical alternative signaling pathway to suppress FAPs adipogenesis.

### Wnt7a maintains cell morphology and induces nuclear localization of Yes-associated protein (YAP)

Actin cytoskeleton disassembly through RhoA-ROCK signaling and subsequent cell shape changes drive adipogenic differentiation of mouse 3T3-L1 preadipocyte cell line and human stromal stem cells (Chen et al., 2018; Nobusue et al., 2014). To determine whether Wnt7a inhibits FAPs adipogenesis through modulation of cell morphology, we expanded freshly isolated FAPs for 4 days and then cultured the cells in either growth and adipogenic differentiation, with or without Wnt7a (**Fig. 3A**). FAPs in the adipogenic condition (ADM) exhibited significantly reduced cell area compared to the growth condition (GM; **Fig. 3B,C**; p<0.0001). Note that this decrease in cell area is also accompanied by elevated levels of nuclear PPARγ (Fig. 3B). In ADM, Wnt7a significantly increased cell area and max feret diameter compared to its Wnt7a-free control (**Fig. 3B-D**; p<0.0001). In the spontaneously differentiating growth condition (GM), Wnt7a also significantly increased both cell area and max ferret diameter compared to its Wnt7a-free control (**Fig. 3B-D**; p<0.001). Wnt7a-treated FAPs in ADM also exhibited cell area and morphology comparable to FAPs maintained in Wnt7a-free GM (**Fig. 3B,C**). Therefore, these results suggest that Wnt7a in the adipogenic condition prevents the shrinking of cell area and maintains morphology. We also note that Wnt7a induced apparent stress fiber formation in GM, but not in ADM (**Fig. 3B**). This observation suggests that Wnt7a may potentially promote fibrogenesis of FAPs in a context-dependent manner.

**Figure 3.**
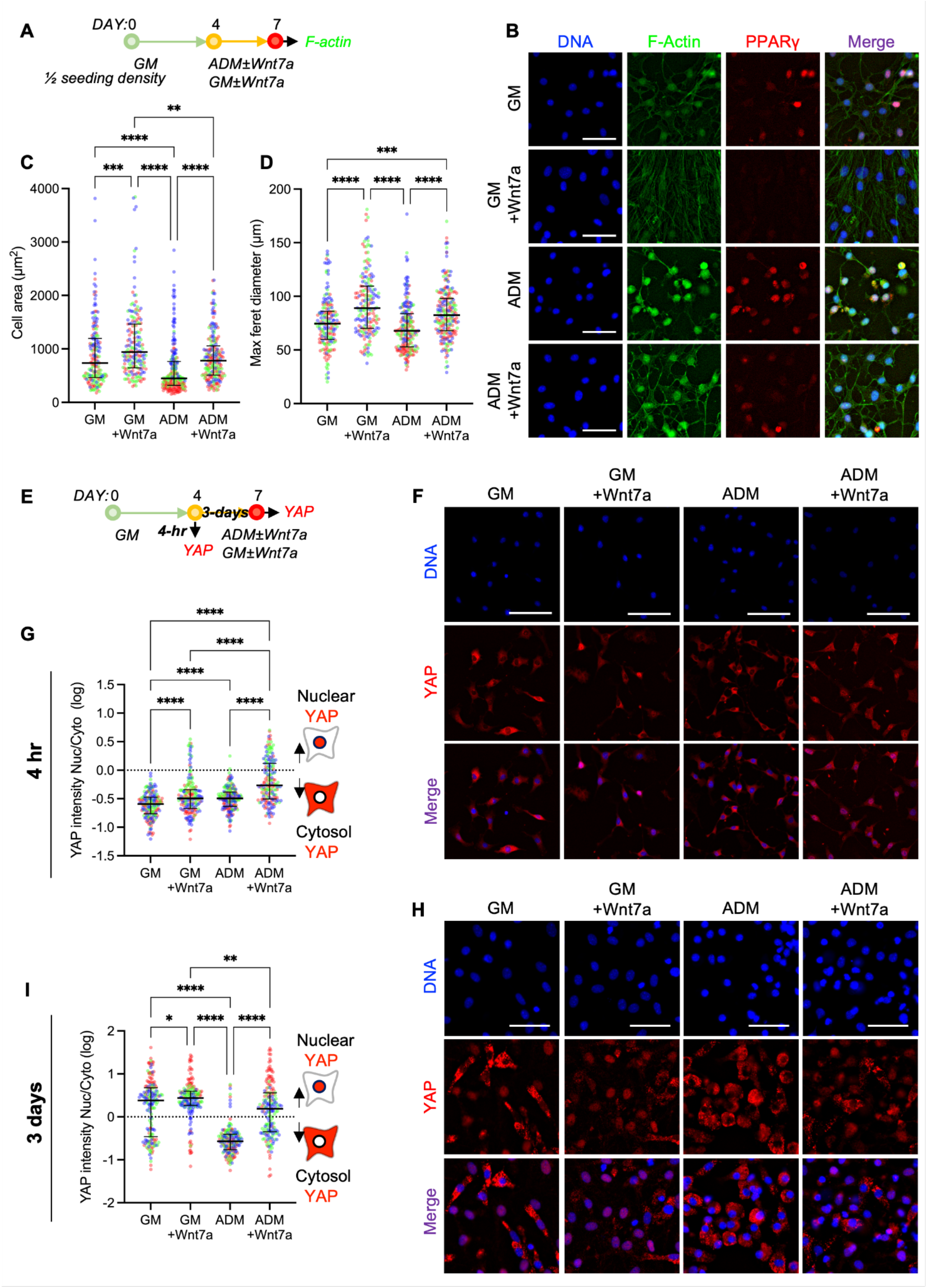
Wnt7a induces nuclear localization of YAP. ***(A)*** Experimental timeline of cell morphology quantification. GM: growth media; ADM: adipogenic differentiation media. ***(B)*** Representative images of F-actin and PPARγ-labeled FAPs. Scale bar: 50 µm. ***(C)*** Cell area quantification. Kruskal-Wallis test with Dunn’s multiple comparisons. Median ± IQR. ** p<0.01; *** p<0.001; **** p<0.0001. n=179-219 cells analyzed from 3 biological replicates. ***(D)*** Max feret diameter quantification. Kruskal-Wallis test with Dunn’s multiple comparisons. Median ± IQR. *** p<0.001; **** p<0.0001. n=179-219 cells analyzed from 3 biological replicates. Colors represent biological replicates ***(C***,***D). (E)*** Experimental timeline of YAP quantification. ***(F)*** Representative immunofluorescence images of YAP-labeled cells after 4-hour treatment in GM ± Wnt7a (200 ng/ml) and ADM ± Wnt7a (200 ng/ml). Scale bar: 100 µm. ***(G)*** Quantification of YAP nuclear:cytosol intensity ratio of 4-hour time point. Values were log-transformed. Kruskal-Wallis with Dunn’s post-hoc analyses. Median ± IQR. **** *p*<0.0001. n=180 cells analyzed from 3 biological replicates. Colors represent biological replicates. ***(H)*** Representative immunofluorescence images of YAP-labeled cells after 3-day treatment in GM ± Wnt7a (200 ng/ml) and ADM ± Wnt7a (200 ng/ml). Scale bars: 25 µm. ***(I)*** Quantification of YAP nuclear:cytosol intensity ratio of 3-day time point. Values were log-transformed. Kruskal-Wallis with Dunn’s post-hoc analyses. Median ± IQR. * *p*<0.05; ** *p*<0.01; **** *p*<0.0001. n=179 cells analyzed from 3 biological replicates. Colors represent biological replicates.

YAP-1 (Yes-associated protein-1) and its paralogue TAZ (transcriptional co-activator with PDZ-binding motif) act as biochemical mechanotransducers that convert mechanical cues and resulting cellular changes (e.g., cell contractility and shape) into cell-specific transcriptional activities (Dupont et al., 2011). Recent evidence also suggests YAP/TAZ also act as downstream modulators of Wnt pathways (Azzolin et al., 2012, 2014; Park et al., 2015a). Based on such evidence and our observation that Wnt7a inhibits FAPs adipogenesis by maintaining cellular shape (**Fig. 3B-D**), we next questioned whether non-canonical Wnt7a signaling promotes nuclear localization of YAP (**Fig. 3E**). Nearly all proliferating FAPs exhibited higher cytosolic YAP in vitro (**Fig. 3F,G**). Culturing the proliferating FAPs in Wnt7a containing growth (GM) and adipogenic (ADM) media for 4 hours significantly increased YAP nuclear-to-cytosolic ratio compared to their respective controls (**Fig. 3F,G**; p<0.0001). YAP nuclear localization was also significantly higher in ADM containing Wnt7a compared to GM containing Wnt7a (**Fig. 3F,G**; p<0.0001), suggesting context-dependent responsivity. These results indicate that Wnt7a promotes nuclear localization of YAP in proliferating FAPs in vitro.

To further determine the effect of prolonged exposure of FAPs to Wnt7a on YAP activity, we cultured FAPs in growth (GM) and adipogenic (ADM) conditions with or without Wnt7a supplementation for 3 days (**Fig. 3E**). In GM, we observed a bimodal distribution of cells expressing nuclear and cytosolic YAP (**Fig. 3H,I**), likely indicating the bifurcating lineage commitment of FAPs, but Wnt7a significantly increased nuclear localization of YAP (**Fig. 3H,I**; p<0.05). By day 3, FAPs undergoing adipogenesis in ADM uniformly exhibited cytosolic YAP (**Fig. 3H,I**). In ADM, 3-day treatment with Wnt7a significantly increased and retained nuclear YAP compared to its control (**Fig. 3H, I**; p<0.0001). In addition, this Wnt7a treatment also resulted in a comparable distribution of cells exhibiting nuclear and cytosolic YAP (i.e., bimodal) with GM without Wnt7a (**Fig. 3H,I**; p>0.05). Collectively, these data suggest that Wnt7a treatment promotes nuclear localization of YAP and continues to retain YAP within the nucleus.

### Wnt7a promotes YAP nuclear localization through Rho

We next asked whether Wnt7a promotes nuclear localization of YAP through the alternative Wnt signaling axis involving Wnt-Frizzled/ROR-Rho GTPases-Lats1/2 (Park et al., 2015a). In the alternative Wnt signaling, inhibition of Rho prevents Wnt-induced YAP activation. To mechanistically test this hypothesis, we pretreated FAPs in adipogenic media (ADM) containing Rho inhibitor (purified C3 Transferase; CT04) or vehicle (dH_2_O) for 2 hours (**Fig. 4A**). The cells were further maintained in ADM with or without CT04 and Wnt7a for additional 4 hours (**Fig. 4A**). Rho inhibition alone did not significantly affect the nuclear localization of YAP compared to the control (**Fig. 4B,C**), but significantly increased the frequency of cells with “diffuse” YAP (**Fig. 4D**). Wnt7a significantly increased YAP nuclear localization compared to the control (**Fig. 4B-D**; p<0.001). However, when Rho was inhibited, Wnt7a failed to promote nuclear localization of YAP (**Fig. 4B,C**; p>0.05 vs control; p<0.0001 vs Wnt7a only). Furthermore, Wnt7a treatment with Rho inhibition resulted in no change in the frequency of FAPs exhibiting cytosolic, diffuse, and nuclear YAP compared to CT04 treatment alone (**Fig. 4D**). In sum, the results suggest that Rho is required for Wnt7a-induced activation of YAP in FAPs.

**Figure 4.**
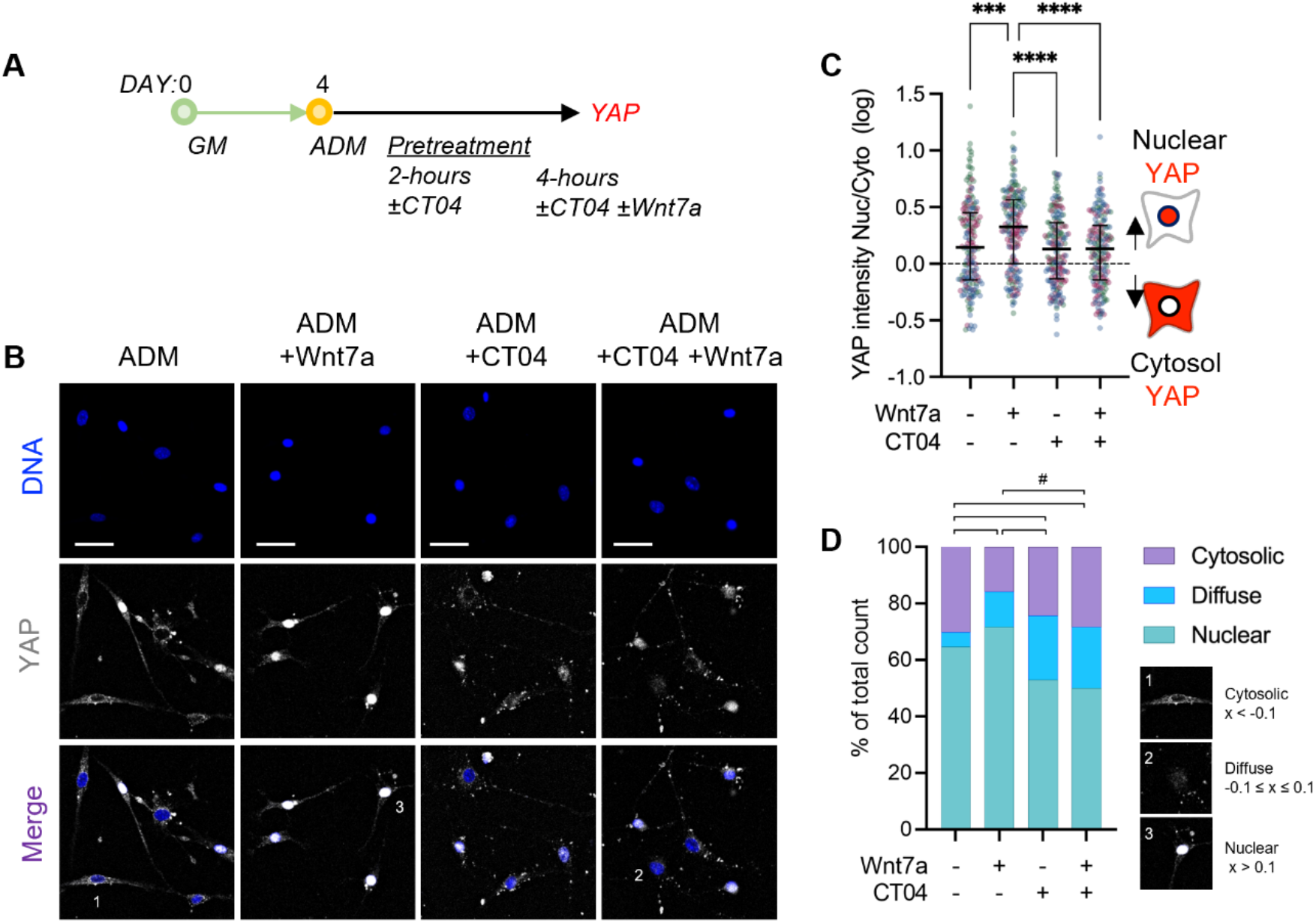
Wnt7a activates YAP through Rho-ROCK and rescues contractility-induced adipogenesis. ***(A)*** Experimental timeline of Rho inhibition study. GM: growth media; ADM: adipogenic differentiation media. ***(B)*** Representative immunofluorescence images of YAP-labeled cells. Scale bar: 50 µm. ***(C)*** Quantification of YAP nuclear:cytosol intensity ratio. Values were log-transformed. Kruskal-Wallis with Dunn’s post-hoc analyses. Median ± IQR. *** *p*<0.001; **** *p*<0.0001. n=180 cells analyzed from 3 biological replicates. Colors represent biological replicates. ***(D)*** Quantification of the cell proportions with nuclear (log(Nuc:Cyto)>0.1), diffuse (−0.1≤log(Nuc:Cyto)≤-0.1), and cytoplasmic (log(Nuc:Cyto)<-0.1) YAP localization. Chi-squared tests with Bonferroni correction. # *p*<0.008.

### Wnt7a promotes downstream canonical signaling through YAP/TAZ nuclear localization

TAZ, a paralogue of YAP-1, regulates the differentiation potential of mesenchymal stem cells by directly repressing PPARγ while activating Runx2 genes (Hong et al., 2005). Furthermore, the repressive role of Wnt/β-catenin in adipogenesis of mesenchymal stem cells also has been well-established (Kang et al., 2007). This is also validated in muscle FAPs in this study, where inhibiting β-catenin activity with PNU74654 increased adipogenesis (**Fig. 2D-E**). Based on such evidence, we hypothesized that non-canonical Wnt7a increases YAP/TAZ activity and subsequently promotes the secretion of modulators of canonical Wnt signaling (e.g., Wnt ligands and Dkk1) to cooperatively inhibit adipogenesis. To test this, we treated proliferating FAPs with Wnt7a for 1 day in culture and immunostained for TAZ. Wnt7a significantly increased nuclear localization of TAZ (**Fig. 5A-C**; p<0.0001), comparable to YAP (**Fig. 3E-H**). Next, to determine if Wnt7a promotes the secretion of modulators of canonical Wnt signaling upon YAP nuclear localization, we treated FAPs treated with Wnt7a for 2 days and performed gene expression analyses on an array of Wnt-related genes (**Fig. 5D**). This time point was chosen because Wnt7a induces YAP nuclear localization without increasing β-catenin nuclear localization at 4 hours post-treatment (**Fig. 2, 3**). Thus, any changes in the canonical Wnt signaling genes after 2-day Wnt7a treatment is likely induced following the YAP nuclear localization. We found that Wnt7a significantly upregulated several canonical Wnt genes including β-catenin, Wisp1, Fzd, and Wnt9a in FAPs (**Fig. 5D-E**; p<0.05). Wnt7a significantly downregulated DKK1 (Dickkopf-related protein 1; p<0.05), which inhibits Lrp5,6 co-receptors that activate the canonical signaling pathway, further supporting our hypothesis. Other upregulated genes include Wnt11, Wnt2, Wnt16, Axin2, Wnt6, and Wnt5a. To further corroborate this, we immunolabeled FAPs with anti-β-catenin 2 days following the Wnt7a treatment. FAPs treated with Wnt7a exhibited significantly increased nuclear β-catenin intensity (**Fig. 5F**; p<0.05). Taken together, these data suggest that Wnt7a first activates YAP/TAZ nuclear localization through Rho (**Fig. 3-5**) and subsequently promotes canonical Wnt signaling (**Fig. 5D,F**) to act in an autocrine manner to inhibit adipogenesis (**Fig. 7A,B**).

**Figure 5.**
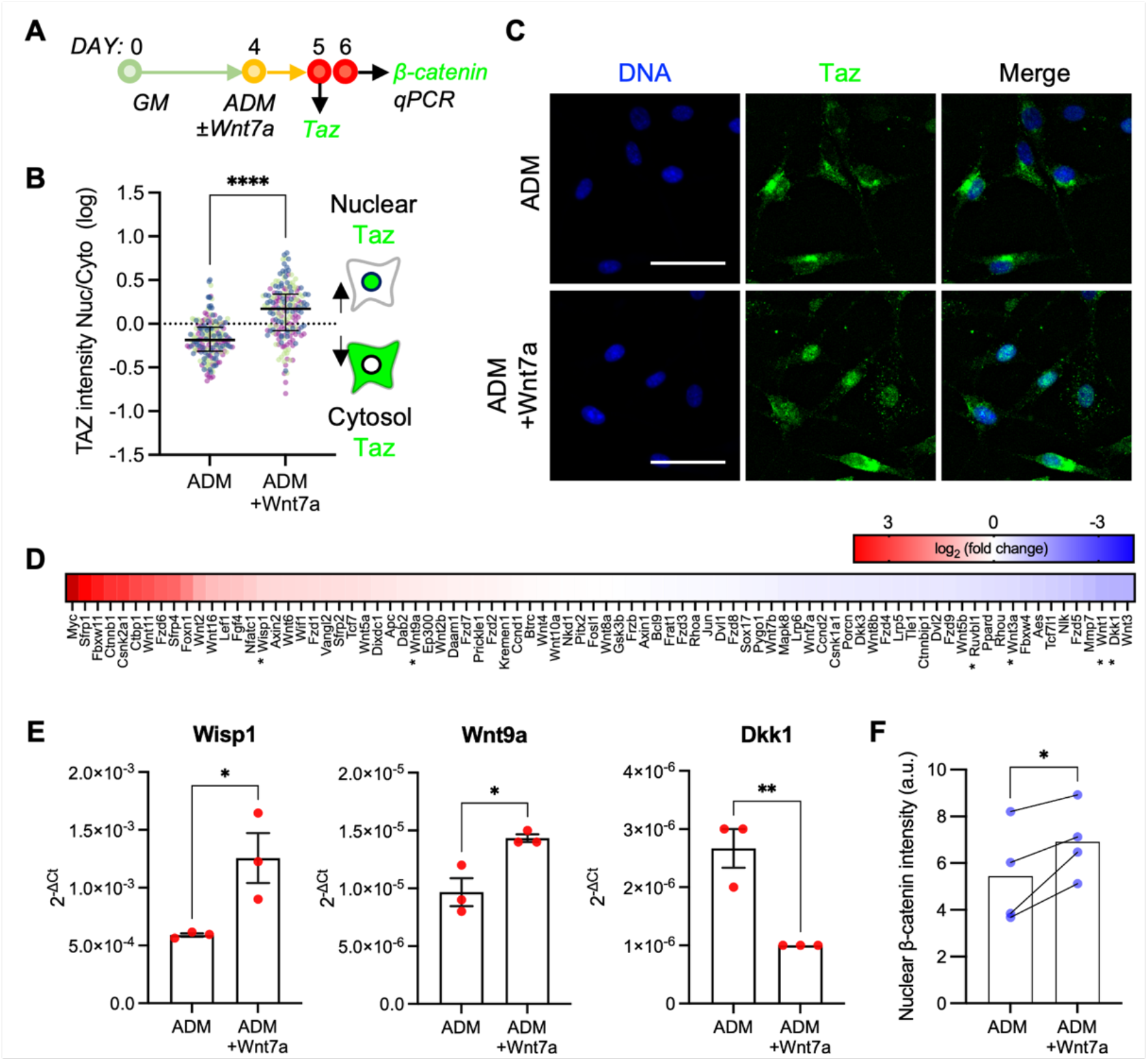
Wnt7a activates TAZ and subsequently activates canonical targets. ***(A)*** Experimental timeline for TAZ, β-catenin, and qPCR. ***(B)*** Quantification of TAZ nuclear:cytosol intensity ratio. Values were log-transformed. Unpaired t-test. **** p<0.0001. n=3. Colors represent biological replicates. ***(C)*** Representative immunofluorescence images of TAZ-labeled cells. Scale bar: 50 µm. ***(D)*** Gene expression analysis heat map array of Wnt-related genes. n=3. * denotes statistically significant differences in fold change. ***(E)*** Relative fold difference of Wisp1, Wnt9a, and DKK1. Unpaired t-test. Mean ± SEM. * p<0.05; ** p<0.01. n=3. ***(F)*** Median nuclear β-catenin intensity (a.u.) after 2 days of Wnt7a treatment. Paired t-test. n=4. * p<0.05.

### Wnt7a suppresses fatty infiltration in skeletal muscle

We next sought to evaluate the efficacy of Wnt7a in suppressing intramuscular fatty infiltration in vivo using the glycerol injury model. Intramuscular injection of glycerol stimulates a robust and reproducible fatty infiltration without affecting muscle regeneration, thus serving as an excellent proof-of-concept in vivo model to evaluate therapeutics targeted for reducing intramuscular adipogenesis (Pisani et al., 2010). To induce injury, tibialis anterior (TA) muscles were injected with glycerol (50% v/v). Wnt7a or saline was injected into the belly of the injured TA muscles 1-day post-injury (**Fig. 6A**). The muscles were then harvested 14-day post-injury for analyses (**Fig. 6A**). Glycerol-injured TAs with saline treatment exhibited a severe fatty infiltration, marked by perilipin-1 expression within the interstitial space (**Fig. 6B; Supp. Fig. 4A**). However, glycerol-injured TAs with Wnt7a treatment exhibited a significantly reduced perilipin-1 expression (**Fig. 6B, C; Supp. Fig. 4A**; p<0.05). We observed no statistically significant differences in the myofiber area distribution (**Fig. 6D, E**). We also observed no qualitative differences in fibrosis assessed by trichrome staining (**Supp. Fig. 4B**). These data collectively show that Wnt7a effectively suppresses intramuscular fatty infiltration in vivo without negatively impacting myogenesis nor inducing fibrosis.

**Figure 6.**
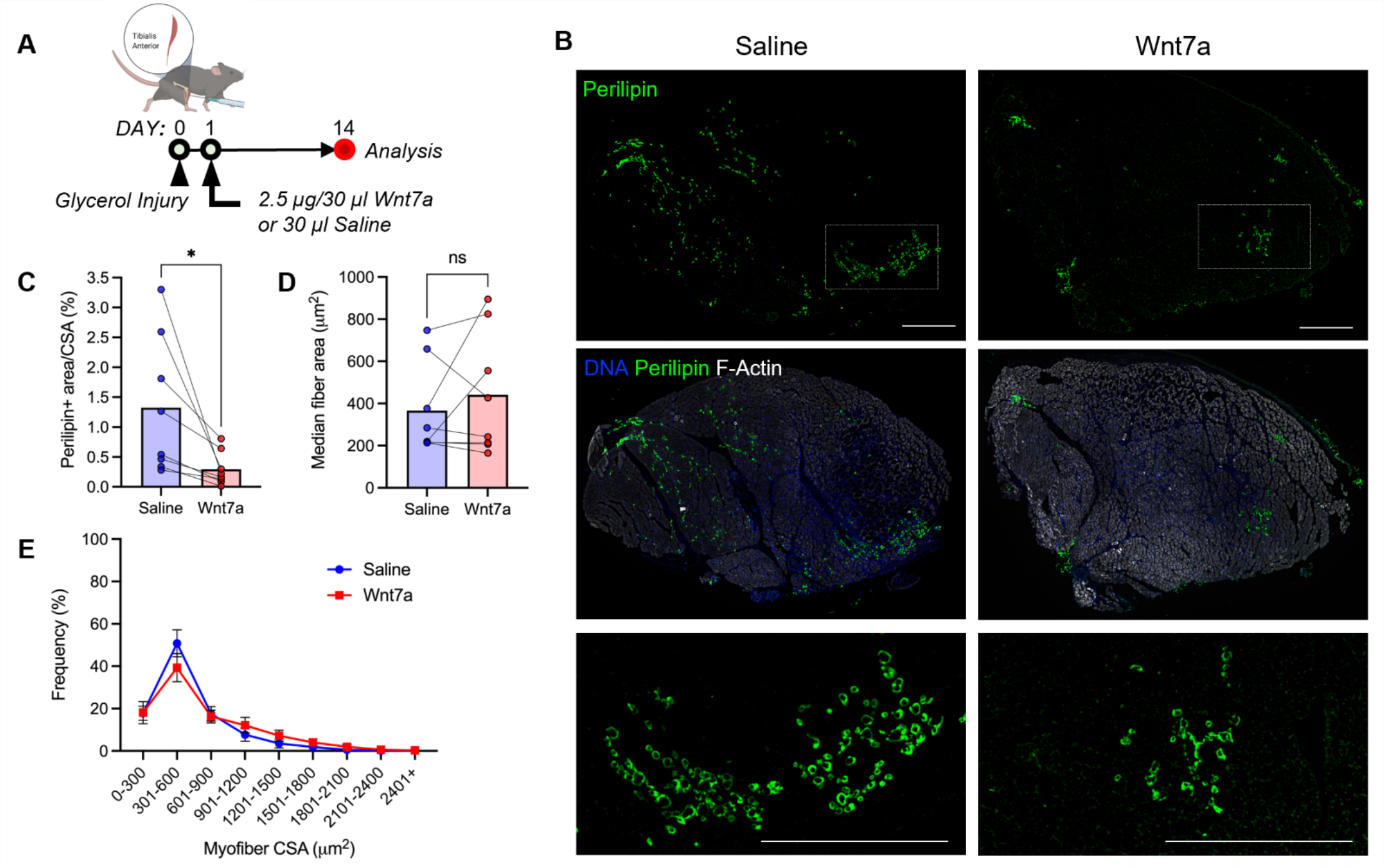
Wnt7a suppresses fatty infiltration in skeletal muscle. ***(A)*** Experimental timeline outlining in vivo glycerol injection, Wnt7a administration, and analyses. Created with Biorender.com. ***(B)*** Representative cross-sections of glycerol-injured TA muscles treated with Wnt7a or saline. Scale bars: 500 µm. ***(C)*** Perilipin area normalized by the TA cross-sectional area. Two-tailed paired t-test. * p<0.05. ***(D)*** Median fiber cross-sectional area. Two-tailed paired t-test. ***(E)*** Histogram of fiber cross-sectional area. Mean ± SEM.

## DISCUSSION

Persistent fatty infiltration is a chronic hallmark of skeletal muscle injuries and diseases, such as rotator cuff injuries (Fu et al., 2021). Wnt7a has been emerging as a potential therapeutic for muscle diseases and injuries due to its pro-regenerative effects on muscle satellite cells and myofibers (Han et al., 2019; Le Grand et al., 2009; von Maltzahn et al., 2011), but its effects on FAPs remain unknown. In this study, we determined the mechanistic effect of Wnt7a on adipogenesis of FAPs, which are the precursors to fatty infiltration in skeletal muscle pathology (Joe et al., 2010; Liu et al., 2016; Uezumi et al., 2010; Wosczyna et al., 2012). Our data reveal that Wnt7a suppresses adipogenesis through the Rho-YAP/TAZ signaling axis that indirectly promote canonical Wnt signaling.

The mechanistic role of canonical Wnt signaling and β-catenin in suppressing adipogenesis is well-established in other cell types (Bennett et al., 2002; Kang et al., 2007; Longo et al., 2004; Moldes et al., 2003; Reggio et al., 2020; Ross et al., 2000). Thus, we first sought to determine if Wnt7a increased nuclear β-catenin through the canonical pathway in FAPs. However, Wnt7a did not increase nuclear β-catenin in FAPs (**Fig. 2A-C**). Furthermore, Wnt7a remained effective in reducing adipogenesis even when β-catenin activity was inhibited using PNU-74654 (**Fig. 2D-E**), suggesting that Wnt7a acts through a non-canonical pathway. We also observed that Wnt7a retained the stellate cellular morphology of FAPs in adipogenic culture conditions while preventing adipogenesis (**Fig. 3A-D**). Based on these findings, we then asked if Wnt7a acts through the alternative Wnt signaling, where YAP/TAZ is activated through the Wnt-Fzd/Ror-Rho GTPases-Lats1/2 signaling axis independent of β-catenin (Park et al., 2015a). In this study, a brief 4-hour Wnt7a treatment induced nuclear localization of YAP in FAPs (**Fig. 3E-G**). We found that Wnt7a failed to induce nuclear localization of YAP when Rho was inhibited using CT04 (**Fig. 4**). We also confirmed that TAZ concurrently is activated with Wnt7a treatment in FAPs (**Fig. 5A-C**). These data thus suggest that Wnt7a activates YAP/TAZ through Rho-dependent alternative Wnt signaling in FAPs. These data also suggest that fatty infiltration that may arise from skeletal muscle unloading (Kaneshige et al., 2022) could be compensated by potentially delivering exogenous Wnt7a.

How does Wnt7a-induced YAP/TAZ nuclear localization inhibit PPARγ and adipogenesis of FAPs? First, published evidence suggests that nuclear activity of YAP/TAZ inhibits adipogenesis by repressing PPARγ in multiple cell types (Deng et al., 2019; El Ouarrat et al., 2020; Hong et al., 2005; Lorthongpanich et al., 2019; Pan et al., 2018). Specifically, TAZ directly represses PPARγ while activating Runx2 genes in mesenchymal stem cells (Hong et al., 2005). The same mechanism may also be inhibiting the adipogenesis of FAPs when treated with Wnt7a. Second, YAP/TAZ activity and TEAD-induced transcription may trigger the secretion of canonical Wnt ligands and inhibitors (Park et al., 2015a). Wnt/β-catenin signaling is a crucial mediator of adipogenesis, where its downregulation results in the differentiation of preadipocytes into mature adipocytes (Bennett et al., 2002; Longo et al., 2004; Ross et al., 2000). In this study, we also observed that inhibiting β-catenin activity using PNU-74654 increased adipogenesis of FAPs (**Fig. 2D, E**), suggesting a potential involvement of canonical Wnt signaling. Here, we report that while Wnt7a does not immediately activate β-catenin unlike the canonical Wnt3a (**Fig. 2A-C**), Wnt7a may indirectly activate the canonical Wnt pathway through YAP/TAZ. Gene expression analyses revealed that some Wnt ligands and their effectors (e.g., β-catenin, Wisp1, Fzd, Wnt9a, Wnt11, Wnt2, Wnt16, Axin2, Wnt6, and Wnt5a) are upregulated while canonical Wnt pathway suppressors (e.g., Dkk1) are downregulated (**Fig. 5D-E**). We further confirmed that nuclear β-catenin immunostaining intensity was also significantly increased after 2 days of Wnt7a stimulation (**Fig. 5F**). Taken together, Wnt7a inhibits adipogenesis of FAPs by non-canonically activating YAP/TAZ which subsequently modulates the canonical Wnt pathway to suppress PPARγ (**Fig. 7B**).

**Figure 7.**
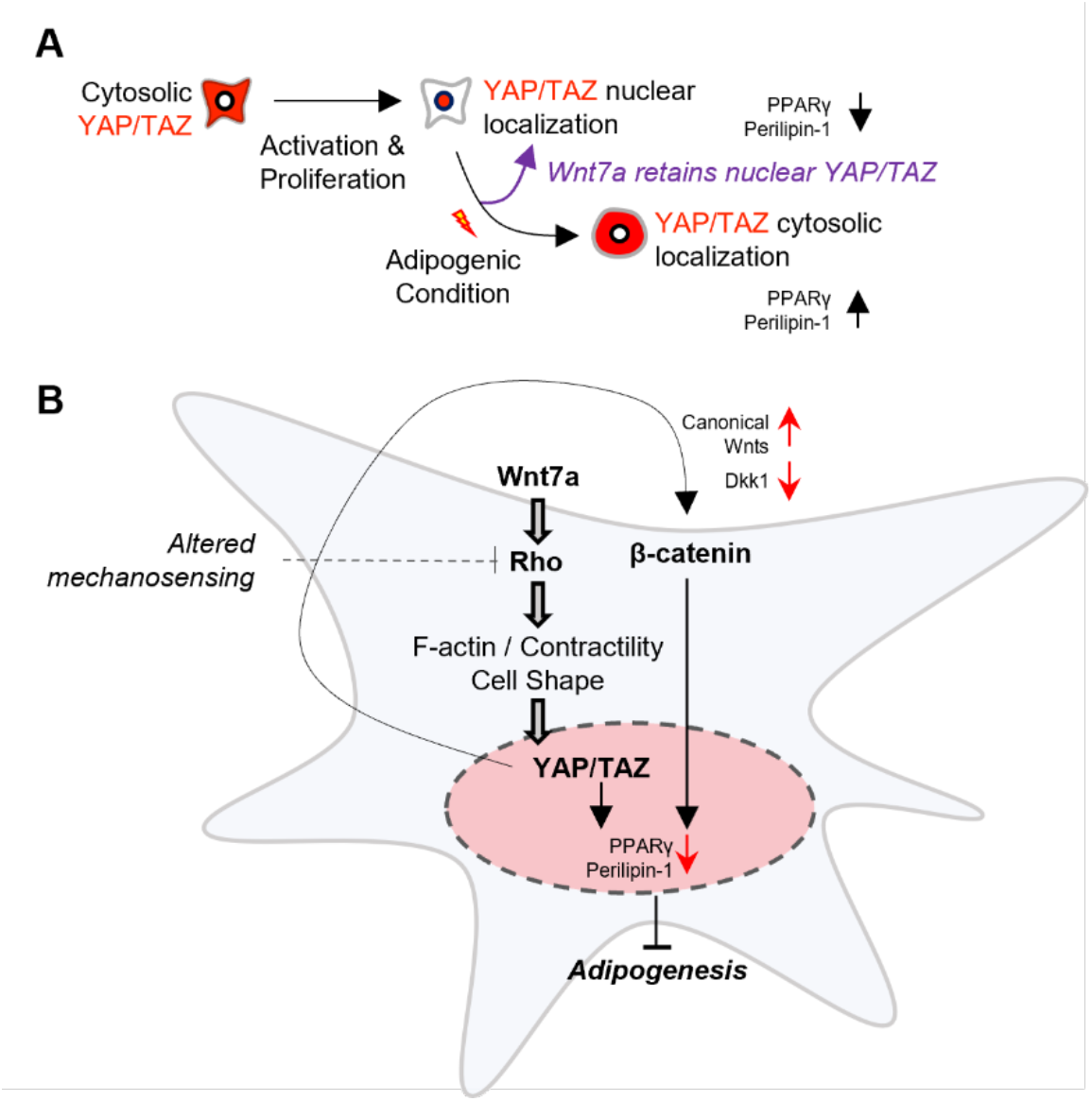
Wnt7a activates YAP/TAZ through Rho and inhibits FAPs adipogenesis. ***(A)*** YAP/TAZ translocates to nucleus in activated and proliferating FAPs. During adipogenic differentiation, YAP/TAZ translocates to cytosol. Wnt7a promotes nuclear translocation and continues to retain YAP/TAZ within the nucleus, thereby preventing FAPs adipogenesis. ***(B)*** Wnt7a may rescue mechanical unloading induced FAPs adipogenesis by reinforcing YAP/TAZ activity through cell contractility mediated mechanisms.

The current study has its limitations. Our current in vitro experiments applied a narrow timeframe in which these FAPs are either spontaneously differentiating or induced to differentiate in cell culture settings. Future investigations will validate the observed mechanisms using clinically relevant in vivo injury and disease models. The cellular identity of the Wnt7a-treated FAPs in vivo is also an important consideration. To further develop Wnt7a as therapeutic, potential crosstalk between FAPs and other cell populations should be considered in vivo in a context-dependent manner as well. Functional measures, including muscle contractile force, mouse gait, and fibrosis, should also be considered to comprehensively evaluate Wnt7a as a potential therapeutic. Ultimately, an effective delivery method using biomaterials such as engineered hydrogels should be tested to show the effectiveness and efficiency of Wnt7a release on therapeutic models.

In conclusion, we identified that Wnt7a inhibits adipogenesis of FAPs through a non-canonical Rho-YAP/TAZ signaling axis. Our data provide insight into applying Wnt7a as a potential therapeutic for mitigating intramuscular fatty infiltration in various skeletal muscle pathologies.

## MATERIALS AND METHODS

### Mice

All animal procedures were conducted under the approved protocol by the Icahn School of Medicine at Mount Sinai Institutional Animal Care and Use Committee. Mice were housed and maintained in the Center for Comparative Medicine and Surgery Facility of the Icahn School of Medicine at Mount Sinai. C57BL/6J mice were acquired from the Jackson Laboratory (stock # 000664). Both male and female mice were used in a randomized manner.

### Glycerol Injuries and Wnt7a Administration

2-month-old mice were anesthetized with isoflurane. 50 µl of 50% glycerol in saline was intramuscularly injected into both tibialis anterior (TA) muscles. After 24 hours, 2.5 µg/30 µl of recombinant human Wnt7a (PeproTech) and 30 µl of saline were injected into the TA muscles in a randomized manner. Buprenorphine shots were subcutaneously administered every 12 hours at the onset of the procedure for 3 days. Mice were sacrificed on day 14 for histological analyses.

### Isolation of Fibro-Adipogenic Progenitors (FAPs)

Primary fibro-adipogenic progenitors (FAPs) were isolated from 4-6-week-old mice by magnetic-activated cell sorting (MACS) as described previously (Marinkovic et al., 2019). Mouse hindlimb muscles were dissected and incubated in digestion media (2.5 U/mL Dispase II; Thermo and 0.2% w/v Collagenase Type II; Worthington in DMEM) on a shaking incubator at 37°C for 1.5 hours. Deactivation media (20% FBS in Ham’s F-10; Gibco) was added to inactivate the reaction. The muscle digest was filtered through a 70 µm cell strainer, then centrifuged (300xg, 5 minutes, 4°C). Cell pellets were resuspended in Staining Buffer (0.5% Bovine Serum Albumin and 2 mM EDTA in PBS) and filtered through a 35 µm cell strainer. Cells were incubated with Biotin anti-mouse CD31 (Biolegend; Cat. No. 102503; 1:150), Biotin anti-mouse CD45 (Biolegend; Cat. No. 103103; 1:150), and Biotin anti-integrin *α*7 (Miltenyi; Cat. No. 130-101-979; 1:10) antibodies at 4°C for 45 minutes. Cells were pelleted through centrifugation and incubated with 10 µl Streptavidin beads (1:30) at 4°C for 15 minutes. Labeled cells were passed through an LD column (Miltenyi) for negative selection. The remaining cells were incubated with Biotin anti-mouse Ly-6A/E (Sca-1) antibody (Biolegend; Cat. No. 122504; 1:75) at 4°C for 20 minutes then 10 µl Streptavidin beads (1:30) at 4°C for 10 minutes. Cells were then passed through an LS column (Miltenyi) and enriched for Sca-1^+^ cells. Cells were filtered through a 35 µm cell strainer once more before cell seeding.

### FAPs Culture and Differentiation

Isolated FAPs were seeded at ∼10,000 cells per 1.0 cm^2^ well in Growth Media (10% FBS and 1x Penicillin-Streptomycin in DMEM) containing 2.5 ng/mL bFGF (Peprotech) on laminin (Gibco; 10 µg/mL) and collagen I (ThermoFisher; 5 µg/mL) coated plates. Cultures were maintained at 37°C and 5% CO_2_ levels. Adipogenic differentiation was performed by incubating the FAPs in Adipogenic Differentiation Media: 0.5 mM 3-isobutyl-1-methylxanthine (MilliporeSigma), 0.25 µM Dexamethasone (MilliporeSigma), and 1 µg/mL insulin (MilliporeSigma) in Growth Media, and Adipogenic Maintenance Media: 1 µg/mL insulin in Growth Media. Fibrogenic differentiation was performed by incubating the FAPs in Fibrogenic Media: 10 ng/mL TGF-β1 (Peprotech) in Growth Media.

### In Vitro Assays and Reagents

Unless otherwise noted, recombinant human Wnt7a (PeproTech; 200 ng/mL; vehicle: dH_2_O) was added to the culture media. To inhibit β-catenin binding to Tcf4, PNU-74654 (Cayman Chemicals; 50 µM; vehicle: DMSO) was added to the media for 3 days plus an additional day in Adipogenic Maintenance Media. To inhibit Rho, CT04 (Cytoskeleton; 2 µg/mL; vehicle: dH_2_O) was added to the media for 2 hours as pre-treatment plus an additional 4 hours with or without Wnt7a.

### Real-Time Quantitative PCR

mRNA was extracted from cells using RNeasy Plus Microkit (Qiagen; Cat. No. 74034) and cDNA was prepared using RT2 First Strand Kit (Qiagen; Cat. No. 330401). RT2 SYBR Green qPCR Master Mix (Qiagen; Cat. No. 330504) and RT2 Profiler PCR Array Mouse WNT Signaling Pathway ABI 7900HT Standard Block plates (Qiagen; Cat. No. PAMM-043ZA) were used for qPCR.

### Immunocytochemistry Staining

Cells were fixed with 4% paraformaldehyde (PFA) for 20 minutes at room temperature. Samples were washed three times with 1x PBS and incubated in blocking/permeabilization buffer (5.0% goat serum, 2.0% bovine serum albumin, 0.5% Triton X-100 in PBS) overnight at 4°C. For PDGFRα staining, blocking/permeabilization buffer without the goat serum (2.0% bovine serum albumin, 0.5% Triton X-100 in PBS) was used. The following primary and secondary antibodies were used for immunocytochemistry in this study: anti-perilipin (Abcam; ab3526; 1:200), anti-alpha smooth muscle actin (Abcam; ab7817; 1:200), anti-YAP (Santa Cruz Biotechnology; sc101199; 1:200), anti-TAZ (Cell Signaling Technology; 83669S; 1:100), anti-PPARγ (Santa Cruz Biotechnology; sc7273; 1:200), anti-PDGFRα (R&D Systems; AF1062; 1:200), anti-β catenin (Cell Signaling Technology; 8480S; 1:100), goat anti-rabbit Alexa-488 (ThermoFisher; A11008; 1:500) and goat anti-mouse Alexa Fluor 546 (ThermoFisher; A11003; 1:500). Hoechst 33342 (ThermoFisher; 1:1000) and Phalloidin-iFluor 488 (Cayman Chemicals; 1:1000) were used to stain nuclei and F-actin, respectively.

### Oil Red O Staining

Cells were fixed with 4% PFA for 20 minutes at room temperature. Samples were washed three times with 1x PBS and incubated in blocking/permeabilization buffer (5.0% goat serum, 2.0% bovine serum albumin, 0.5% Triton X-100 in PBS) overnight at 4°C. Cells were incubated in isopropanol (60%) for 5 min then incubated in Oil Red O for 20 min. Cells were washed five times with 1x PBS and then counterstained with Hoechst 33342 (ThermoFisher; 1:1000).

### Histology and Immunohistochemistry

Hindlimbs were dissected and fixed for 1 hour at room temperature in 4% paraformaldehyde in PBS. Fixed TA muscles were dissected and frozen in liquid nitrogen chilled isopentane. 10 µm sections were obtained from the frozen TAs using a cryotome. Tissue sections were incubated using blocking/permeabilization buffer (5.0% goat serum, 2.0% bovine serum albumin, 0.5% Triton X-100 in PBS) for 1 hour at room temperature. The following primary and secondary antibodies were used for tissue immunohistochemistry in this study: anti-perilipin (Abcam; ab3526; 1:200) and goat anti-rabbit Alexa Fluor 488 (ThermoFisher; A11008; 1:500). Hoechst 33342 (ThermoFisher; PI62249; 1:1000) and Alexa Fluor 647 Phalloidin (ThermoFisher; A22287; 1:1000) were used to stain nuclei and F-actin, respectively. Tissue sections were also processed for routine H&E and trichrome staining.

### Imaging and Image Analysis

Images were taken on a Leica Microsystems THUNDER DMi8 microscope using LAS-X software for processing. Unless noted below, all images were analyzed using ImageJ in an automated manner. YAP/TAZ nuclear-cytoplasmic ratio and cell shape were manually quantified.

### Statistical Analysis

Statistical analyses were performed using GraphPad Prism software. Normality was determined using the Shapiro-Wilk test and Q-Q plot. To test statistical significance, unpaired t-test, one-way analysis of variance (ANOVA) with Tukey’s post-hoc analysis, two-way ANOVA with Bonferroni post-hoc analysis, and Kruskal-Wallis test with Dunn’s multiple comparisons were performed depending on data normality and number of comparisons. p<0.05 was considered statistically significant. All experiments and studies had at least three biological replicates.

## FUNDING

This study was supported by the Department of Orthopaedics at the Icahn School of Medicine at Mount Sinai to Woojin M. Han.

## AUTHOR CONTRIBUTIONS

CF and WMH conceived and designed the studies. CF, MGS, DL, and WMH conducted experiments and analyzed data. CF and WMH wrote and revised this manuscript.

## CONFLICTS OF INTERESTS

All authors have no competing interests to disclose.

## DATA AND MATERIALS AVAILABILITY

All data needed to evaluate the conclusion of the paper are present in the paper or the Supplementary Materials.

## ACKNOWLEDGEMENTS

We thank Nada Marjanovic (Sinai qPCR Core) for her work on the qPCR and gene expression analysis.

## Notes

### Competing Interest Statement

The authors have declared no competing interest.

