## Supplementary Materials for "Wnt7a Suppresses Adipogenesis of Skeletal Muscle Mesenchymal Stem Cells and Fatty Infiltration Through the Alternative Wnt-Rho-YAP/TAZ Signaling Axis"

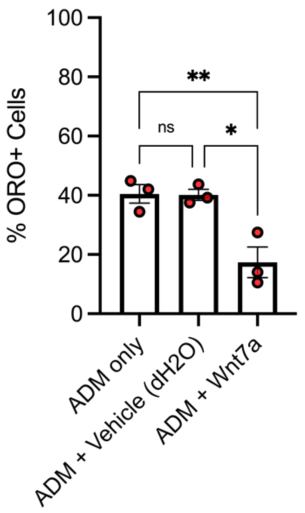


**Supplemental Figure 1.** d$H_2O$ vehicle in ADM (0.2% v/v) does not affect percent ORO+ cells compared to the ADM only condition. One-way ANOVA with Tukey's post-hoc analysis. * $p<0.05$; ** $p<0.01$. n=3.

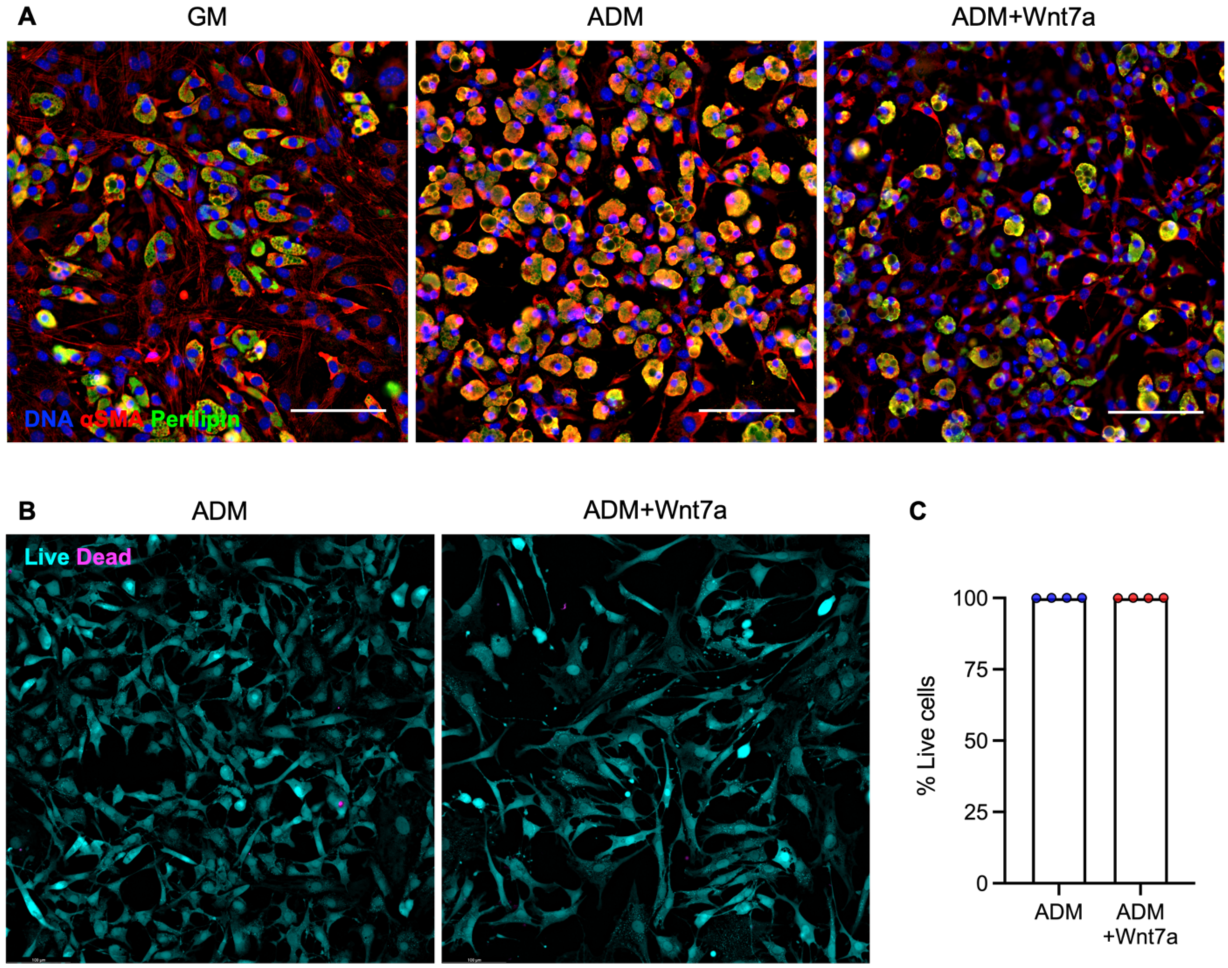


**Supplemental Figure 2.** ***(A)*** Representative immunofluorescence images of α-smooth muscle actin (αSMA) and perilipin-labeled FAPs. Scale bar: 100 μm. ***(B)*** Representative images of live-dead staining in ADM ± Wnt7a conditions. ***(C)*** Quantification of % live cells show minimal cell death with Wnt7a (200 ng/ml). n=4. Scale bar: 100 μm.

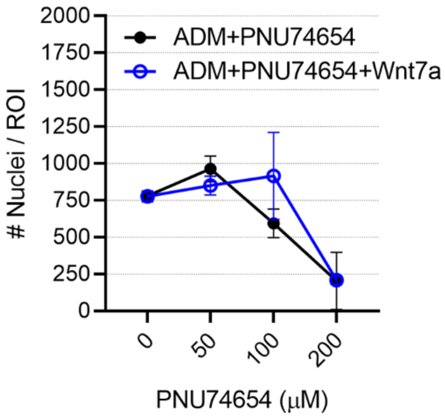


**Supplemental Figure 3.** Dose-response assay of PNU74654. Nuclei count begins to decrease around 100 μM after 5 days. n=2 per concentration.

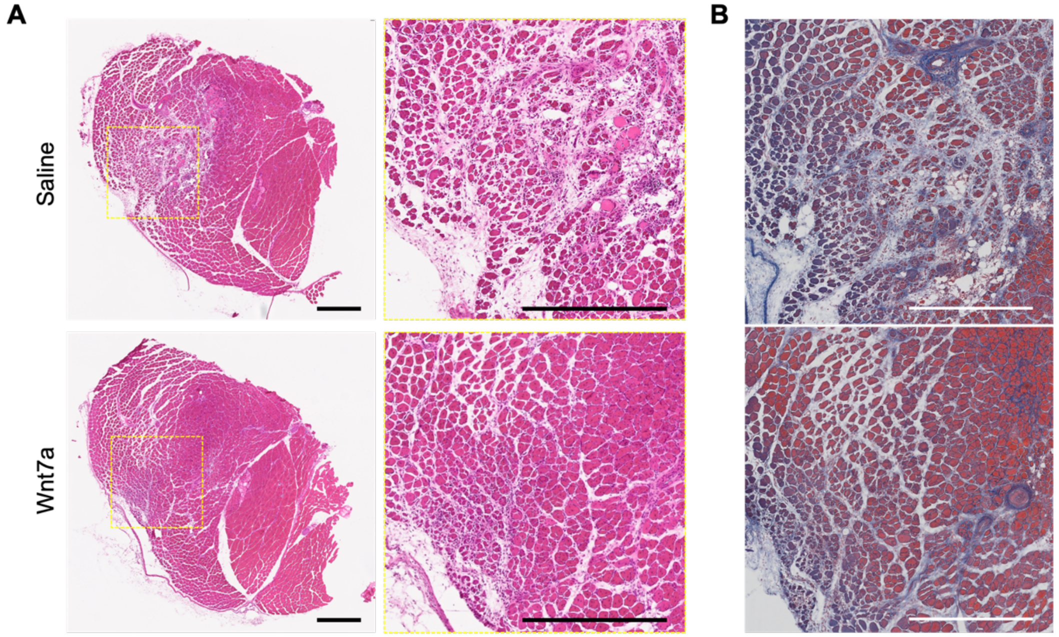


**Supplemental Figure 4.** ***(A)*** Representative H&E staining of cross-sectioned TA muscles injected with saline (30 μl) ± Wnt7a (2 μg/30 μl) following glycerol-induced injury. Day 14 post-injury. Scale bar: 500 μm. ***(B)*** Representative trichrome staining of cross-sectioned TA muscles injected with saline (30 μl) ± Wnt7a (2 μg/30 μl) following glycerol-induced injury. Day 14 post-injury. Scale bar: 500 μm.
